# Comparison of Type-1 and Type-2 Fuzzy Systems for Mineralization of Bioprinted Bone

**DOI:** 10.1101/2021.03.31.437908

**Authors:** Ashkan Sedigh, Mohammad-R. Akbarzadeh-T, Ryan E. Tomlinson

**Author notes:** Address correspondence to: Ryan Tomlinson, PhD, 1015 Walnut Street, Department of Orthopaedic Surgery, Thomas Jefferson University, Philadelphia, PA, 19107, USA.

## Abstract

Bioprinting is an emerging tissue engineering method used to generate cell-laden scaffolds with high spatial resolution. Bioprinting parameters, such as pressure, nozzle size, and speed, have a large influence on the quality of the bioprinted construct. Moreover, cell suspension density, cell culture period, and other critical biological parameters directly impact the biological function of the final product. Therefore, an approximation model that can be used to find the values of bioprinting parameters that will result in optimal bioprinted constructs is highly desired. Here, we propose type-1 and type-2 fuzzy systems to handle the uncertainty and imprecision in optimizing the input values. Specifically, we focus on the biological parameters, such as culture period, that can be used to maximize the output value (mineralization volume). To achieve a more accurate approximation, we have compared a type-2 fuzzy system with a type-1 fuzzy system using two levels of uncertainty. We hypothesized that type-2 fuzzy systems may be preferred in biological systems, due to the inherent vagueness and imprecision of the input data. Here, our results demonstrate that the type-2 fuzzy system with a high uncertainty boundary (30%) is superior to type-1 and type-2 with low uncertainty boundary fuzzy systems in the overall output approximation error for bone bioprinting inputs.

## 1. Introduction

Additive manufacturing is a process by which three-dimensional objects are generated by material deposition in sequential layers [1][2]. Bioprinting is an emerging field of additive manufacturing, in which bioactive scaffolds can be quickly generated by deposition of layers of cell-laden biocompatible materials, such as collagen or other hydrogels. After specifying the exact geometry of the construct, G-code containing the extrusion path and parameters is generated to direct fabrication by one of several commercially available desktop bioprinters. Indeed, the ability to place cells in biologically relevant scaffold materials with high spatial resolution has made bioprinting a popular fabrication method for tissue engineering [3][4].

In addition to the specific geometry of the bioprinted construct, the parameters used to perform the bioprinting procedure itself will have significant effects on the final properties of the model. Therefore, it is essential to fully characterize and optimize the bioprinting parameters (e.g. print speed or bioink viscosity) that are necessary to reach the desired outputs, such as high cell viability, appropriate cell function, and necessary mechanical properties [5]. For example, increasing the nozzle size on the bioprinter decreases the shear stress placed on the biomaterial during extrusion, resulting in both increased cell viability and reduced print resolution [6]. Therefore, determining the optimal print parameters is important for success in bioprinting. As a result, several studies have been performed in the field of bioprinting optimization, such as optimization of a solid model for 3D bioprinting, bioink optimization, and bioprinting parameter selection [7][8].

Nonetheless, there is a significant degree of imprecision and uncertainty inherent in bioprinting optimization. A potential approach to handling this issue, which arises from normal biological variation, is through the implementation of approximation systems based on computational methods [9]. In recent years, systems biology has become a critical multidisciplinary research area between computer science and biology. Studies in this field aim to develop computational models of biological processes, requiring both a robust dynamic model as well as a large dataset of experimental results [10][11].

Developing a dynamic model is challenging and requires well-characterized control parameters in order to approximate outcomes from laboratory experiments. Nonetheless, recent studies have observed that computational optimization algorithms can effectively approximate output parameters using either a deterministic or stochastic biological model. A partial list of the approaches employed to this end include meta-heuristic, evolutionary, global optimization, genetic programming, simulated annealing, simplex, ant-colony, fuzzy genetic hybrid system, and multi-objective optimization [12][10].

Here, we have developed a quantitative model of a biological system using the fuzzy system approach, which is a potential solution for overcoming uncertainty in an experimental dataset. In fact, previous studies have shown that the accuracy of a fuzzy system approach is the same as the deterministic mathematical approach (ordinary differential equations) for the same kinetic dataset [13]. Moreover, fuzzy systems can be utilized to find the qualitative system response when a quantitative dataset is not available [14][15].

Fuzzy logic is an extended model of standard logic. In standard logic, values can only be completely false or completely true (degree of truth equal to 0 and 1, respectively), whereas fuzzy logic values can have a degree of truth between 0 and 1. This generalization provides a mathematical model to move from discrete to continuous values. A system based on fuzzy logic is called a fuzzy system [16]. In order to define a fuzzy system, it is essential to first describe the *fuzzy set theory*.

In contrast to sets in classical logic, *a fuzzy set* is a set without a crisp boundary. For instance, if the reference set *X* is a Universe of **discourse** for elements *x*, the fuzzy set *A* is defined:

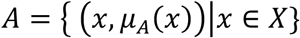

where *μ_A_*(*x*) is called the *Membership Function (MF)* for the fuzzy set *A*. The *MF* maps each element of the Universe set *X* to a grade between 0 and 1. A membership with grade 0 means that the associated element is not included, whereas the membership value of 1 means a fully included element [17]. One of the most important applications of fuzzy set theory, introduced by Zadeh [16], is the fuzzy *rule-based* system. This tool is based on “*if-then*” rules where the antecedents and consequences are fuzzy logic statements. This rule-based fuzzy system is used for modelling the inputs and their relationships with the output variables. A type-1 fuzzy system (T1 FS) is a framework consisting of weighted *rules, membership functions*, and a *fuzzy inference system*. This system takes the crisp data (fuzzy singletons) or fuzzy inputs and generates fuzzy outputs based on the given *if-then rules*. A method of *defuzzification* is then used to extract a crisp value inferred from the fuzzy model [18].

The concept of a Type-2 Fuzzy System (T2 FS) is an extension of an ordinary type-1 fuzzy set which was introduced by Zadeh [19]. In contrast to a T1 FS, Type-2 fuzzy sets have grades of membership that are themselves fuzzy both in *primary and secondary memberships*. A *primary membership* is the same as type-1 membership that maps each element to a grade between 0 and 1. Relative to each primary membership, there is a *secondary membership* (a grade between 0 and 1) that defines the uncertainty in defining the primary membership using a fuzzy set construct. A type-2 FLS is also a framework consisting of weighted *rules (IF-THEN), membership functions*, and a *fuzzy inference system*.

The type-2 fuzzy inference system is also similar to its type-1 counterpart, but includes a type-reducer and defuzzifier, which generate a type-1 fuzzy set output and a crisp number, respectively [20].

T2 FS have been widely applied to a variety of problems where handling uncertainty is critical, including decision making, function approximation, and data preprocessing [21]–[25]. One example is a model with noisy and uncertain training data – here, uncertainty exists in the antecedent and consequents. The level of uncertainty and information regarding it can be used in mathematical modeling of antecedents and consequents. Moreover, non-quantitative data is often disseminated using words that convey an indistinct level of certainty [26]. T2 FS has the capability to grade these linguistic representations into membership functions, which shows a more robust algorithm rather than T1 FS in inferencing input data. However, the computational cost of T2 FS is higher than the T1 FS [20].

The main objective of this study is to implement fuzzy systems towards the optimization of bioprinting parameters. We hypothesize that the implementation of a T2 FS would reduce the error in the output in biological systems with inherent uncertainty in both the inputs and outputs. To directly test this hypothesis, we have implemented type-1 and type-2 fuzzy logic systems and compared the performance of each for use in the optimization of bone bioprinting parameters.

## 2. Type-2 Fuzzy System

The general flowchart of a T2 Mamdani fuzzy logic system is shown in Figure 1a, including the fuzzification, fuzzy inference, type reduction, and defuzzification steps. Considering the crisp inputs (n inputs) and one output:

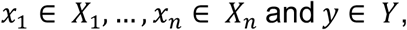

the *k*-th (*k = 1, …, K*) rule in Mamdani T2 FS is expressed as below:

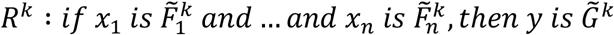

**Figure 1.**
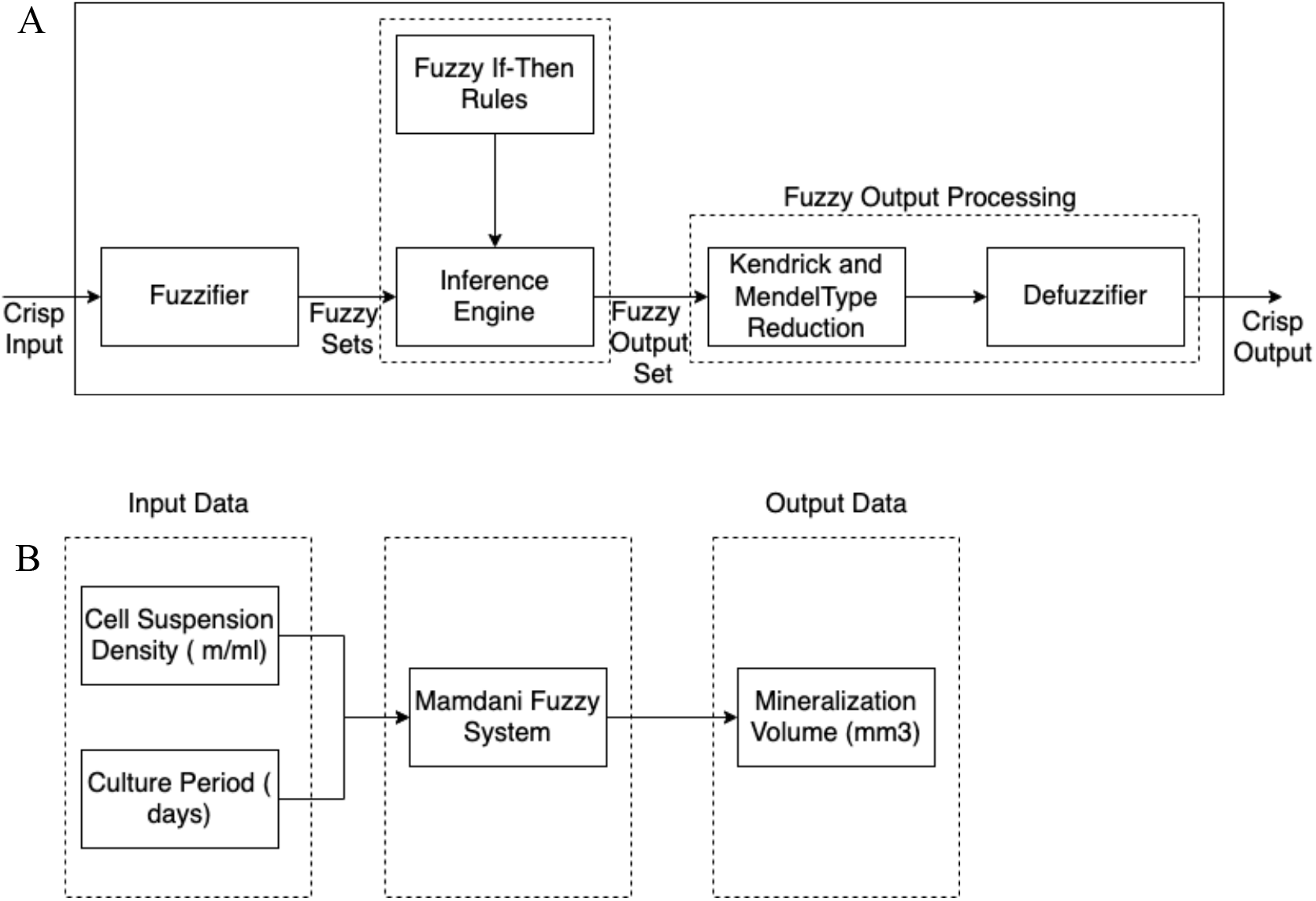
Type-2 Fuzzy Logic Algorithm and Study Design. A) General overview and different features of the Type-2 Fuzzy system including fuzzification, rules, inference engine, type reduction, and defuzzification. B) Schematic of our study design, including two inputs (cell suspension density and culture period) and one output (mineralization volume).

Where 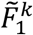 and 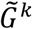 are T2 FS. In this system, rules represent the fuzzy relations between multiple dimensional input space 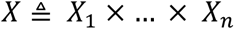 and output space *Y*. The definitions below are paraphrased from the Mendel and Liang article in T2 FS [20]:

#### Definition 1 (Footprint of Uncertainty of a Type-2 Membership Function)

Uncertainty in the region of upper and lower boundary of membership function is called the *footprint of uncertainty*. It is the union of all primary membership grades.

#### Definition 2 (Upper and Lower MFs)

A type-1 fuzzy upper and lower boundary MFs for the Footprint of Uncertainty (FOU) of an interval type-2 MF. The *upper* and *lower* bounds of the region are the maximum and minimum membership grades of FOU.

The over and under bars show the upper and lower MFs, respectively. The membership function of 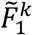 is represented as below:

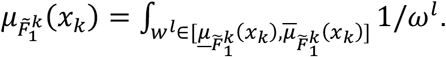

The upper and lower boundaries on a Gaussian primary function with an uncertain standard deviation is represented below. In this representation, the Gaussian primary MF has a fixed mean 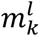 and an uncertain standard deviation 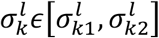:

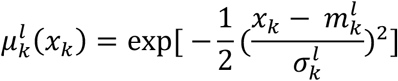

Where

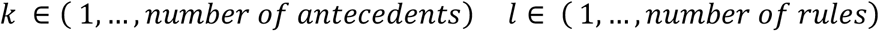

The upper and lower MF of 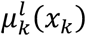 are:

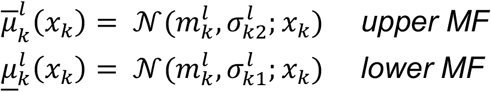

In the interval T2 non-singleton fuzzy system with type-2 fuzzification and minimum or product *t-norm, the output fuzzy set* is represented as below:

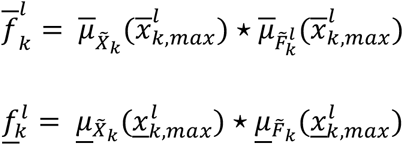

In Equation 2, 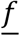 and 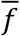 are the result of the input and antecedent operations, based on the value of *x_k_* inwhich the supremum occurs as 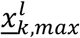 and 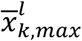:

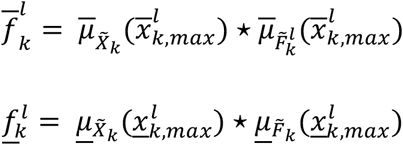

In the algorithm above, we follow the following steps to obtain the crisp output: 1) fuzzification 2) fuzzy inference 3) type-reduction 4) defuzzification. The result of crisp input and antecedents is an interval type-1 fuzzy set, defined by the lower and upper MF 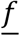 and 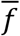, respectively. Regarding Equation (2), the fired output value, which is the combined output consequent set 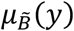, can be computed. The type-2 fuzzy system is computationally intensive to implement [20]. A potential solution is a type-reduction method, such as Kendrick and Mendel proposed using the type-1 defuzzification method for reducing the type-2 to type-1 fuzzy system [27].

Various type-reduction are proposed as centroid, center-of-sets, and height [20]. In our method, we have used center-of-set type-reduction method:

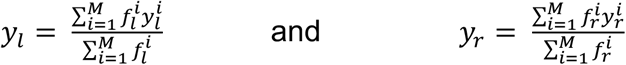

Where the maximum and minimum value of *y* are *y_r_* and y_l_ respectively. *y_r_* and *y_l_* depend only on a mixture of 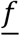 and 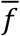 values, because 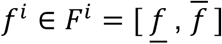. Because *y* is an interval non-convex set, we defuzzify it by using the average of *y_r_* and *y_l_*.

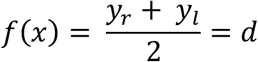

Where *d* is the defuzzified output in the above formula.

## 3. Methods

### A. Fuzzy Inference Engine

The first step to develop this approximation model is to make a fuzzy inference system, such as the Mamdani or Sugeno systems. For this study, we chose the Mamdani inference engine due to the intuitive interpretable nature of its rule-base inferencing [28].

The main difference between Mamdani-type and Sugeno-type Fuzzy Inference System (FIS) is the method of how the output result is obtained. In Mamdani FIS, the crisp result is obtained through defuzzification of the rules. However, Sugeno FIS uses a weighted average of the rules to compute the crisp output. Moreover, Mamdani FIS can be applied to both “Multiple Input, Single Output” (MISO) and “Multiple Input, Multiple Output” (MIMO) systems, which is advantageous for biologic systems that frequently have multiple outputs. It is important to note that Sugeno-type systems can be used for MISO systems [28].

### B. Membership functions parameterization and variables

As discussed earlier, we obtained optimization data for the variables in our model from a previously published experimental study [29]. As shown in Figure 1b, the bioprinting approximation system has two inputs, which are the number of cells (millions per milliliter) and the culture period (days), and one output variable, which is the mineralized volume of the bioprinted construct (mm^3^).

Next, the membership functions (MFs) were defined. MFs are distributed evenly by dividing the full input data range by the number of MFs, which is the input labels shown in Table 1 and Figure 2. The membership function is a 2D curve (type-1 fuzzy) that describes the variables’ degree of membership to a fuzzy set, using a value between 0 and 1. Membership functions are used in a fuzzification process to convert the crisp values to fuzzy values [30]. Membership functions can be implemented using a variety of functions, such as triangular, Gaussian, or gamma. Here, we chose the Gaussian membership function due to the similarity of this function with many biological processes. In general, Gaussian membership functions are popular because of their smoothness and concise notation [31]. MFs designed with Gaussian forms, are modified by tuning the standard deviation and their mean values, as shown in Table 2. Table 1 indicates the fact that low mineralization volume and close data points in 7 days culturing period compared to the 14- and 21-days results in a smaller standard deviation than the two other MFs.

**Table 1.** Input data extracted from the previous study [29]

| Culture Period (days) | Cell Suspension Density<br>(million cells /milliliter) | Mineralization Volume (mm3) |
| --- | --- | --- |
| 7 | 0 | 0 |
| 7 | 1.67 | 0.1 |
| 7 | 5 | 0.1 |
| 7 | 15 | 0.2 |
| 14 | 0 | 0 |
| 14 | 1.67 | 1 |
| 14 | 5 | 3 |
| 14 | 15 | 12 |
| 21 | 0 | 4 |
| 21 | 1.67 | 14 |
| 21 | 5 | 21 |
| 21 | 15 | 24 |

**Figure 2.**
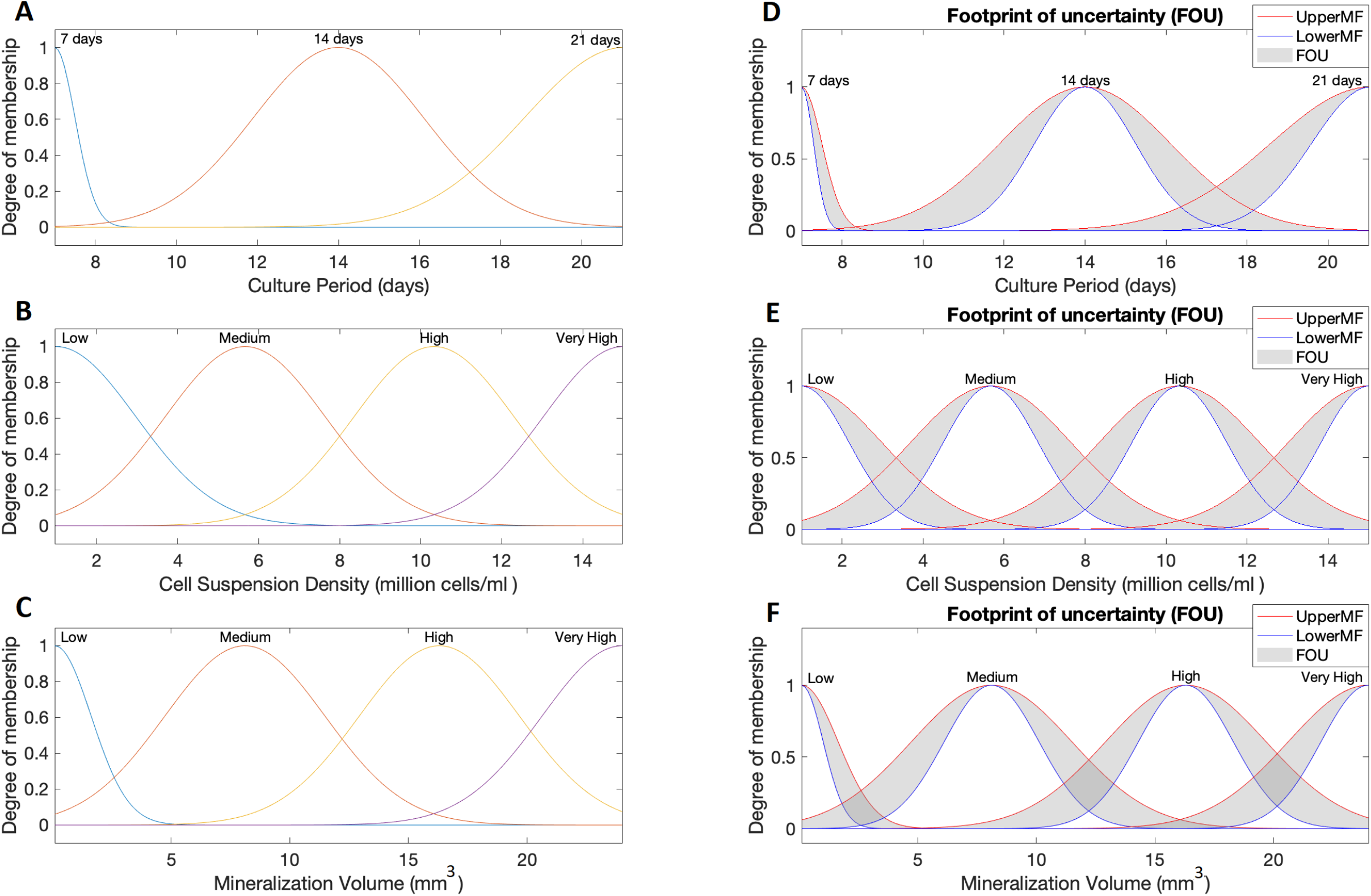
Type-1 and Type-2 Fuzzy Membership Functions. A-C) Type-1 MFs for the two inputs, A) culture period and B) cell suspension density, as well as the single output C) mineralization volume. D-F) Type-2 MFs for D) culture period, E) cell suspension density, and F) mineralization volume. Type-2 membership functions are designed with 0.2 lag. Upper (red) and lower (blue) boundaries are as illustrated.

The membership functions based on the input value representation (Table 1) are plotted in Figure 2 (A, B, C). The Type-1 membership functions were converted to Type-2 with a prescribed 20 and 30 percent uncertainty using the Fuzzy Logic System toolbox (Matlab). The lower and upper boundary of the MFs type-2 FS with 0.2 lag are plotted in Figure 2 (D,E,F).

### C. If-Then Rules Establishment

The next step in designing a fuzzy system is to define the fuzzy IF-THEN rules. As shown in Table 2, we utilized previously published results from an experimental bioprinting study [29]. The model rules are shown at Table 3.

**Table 2.**
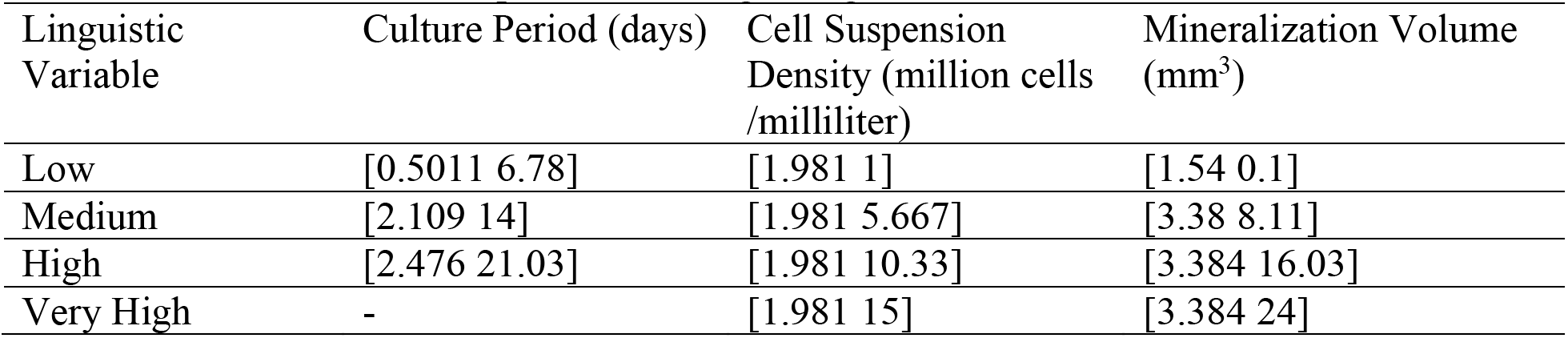
Gaussian membership functions range [Sigma, Mean]

| Linguistic Variable | Culture Period (days) | Cell Suspension Density (million cells /milliliter) | Mineralization Volume (mm<sup>3</sup>) |
| --- | --- | --- | --- |
| Low | [0.5011 6.78] | [1.981 1] | [1.54 0.1] |
| Medium | [2.109 14] | [1.981 5.667] | [3.38 8.11] |
| High | [2.476 21.03] | [1.981 10.33] | [3.384 16.03] |
| Very High | - | [1.981 15] | [3.384 24] |

**Table 3.** Type-1 and Type-2 Fuzzy system rules

| Rule | Culture Period (days) | Cell Suspension Density<br>(million cells / milliliter) | Mineralization Volume (mm<sup>3</sup>) |
| --- | --- | --- | --- |
| 1 | Low | Low | Low |
| 2 | Low | Medium | Low |
| 3 | Low | High | Low |
| 4 | Low | Very High | Low |
| 5 | Medium | Low | Low |
| 6 | Medium | Medium | Medium |
| 7 | Medium | High | Medium |
| 8 | Medium | Very High | High |
| 9 | High | Low | Medium |
| 10 | High | Medium | High |
| 11 | High | High | Very High |
| 12 | High | Very High | Very High |

### D. Fuzzy Inference Process

Finally, we implemented the type-1 fuzzy inference process using the following procedure:

1. Fuzzification of the input variables
2. Application of the fuzzy operator (AND) in the antecedent
3. Implication from the antecedent to the consequent
4. Aggregation of the consequent across the rules
5. Defuzzification

Next, the type-2 fuzzy system is implemented using the following process:

1. Fuzzification of the input variables
2. Application of the fuzzy operator (AND) in the antecedent
3. Convert T1 MF to T2 MF with 0.2 and 0.3 lag
4. Implication from the antecedent to the consequent
5. Aggregation of the consequent across the rules
6. T2 FS to T1 FS type reduction by Karnik-Mendel
7. Defuzzification

In the above process, we used “Minimum” for AND (Step 2), “Minimum” for Implication (Step 4), “Maximum” for Aggregation (Step 5), and “Centroid” for defuzzification (Step 7). The 3D surface of the implemented type-1 fuzzy and type-2 fuzzy (20% and 30% uncertainty) rule based on the two inputs and one output is plotted in Figure 3.

**Figure 3.**
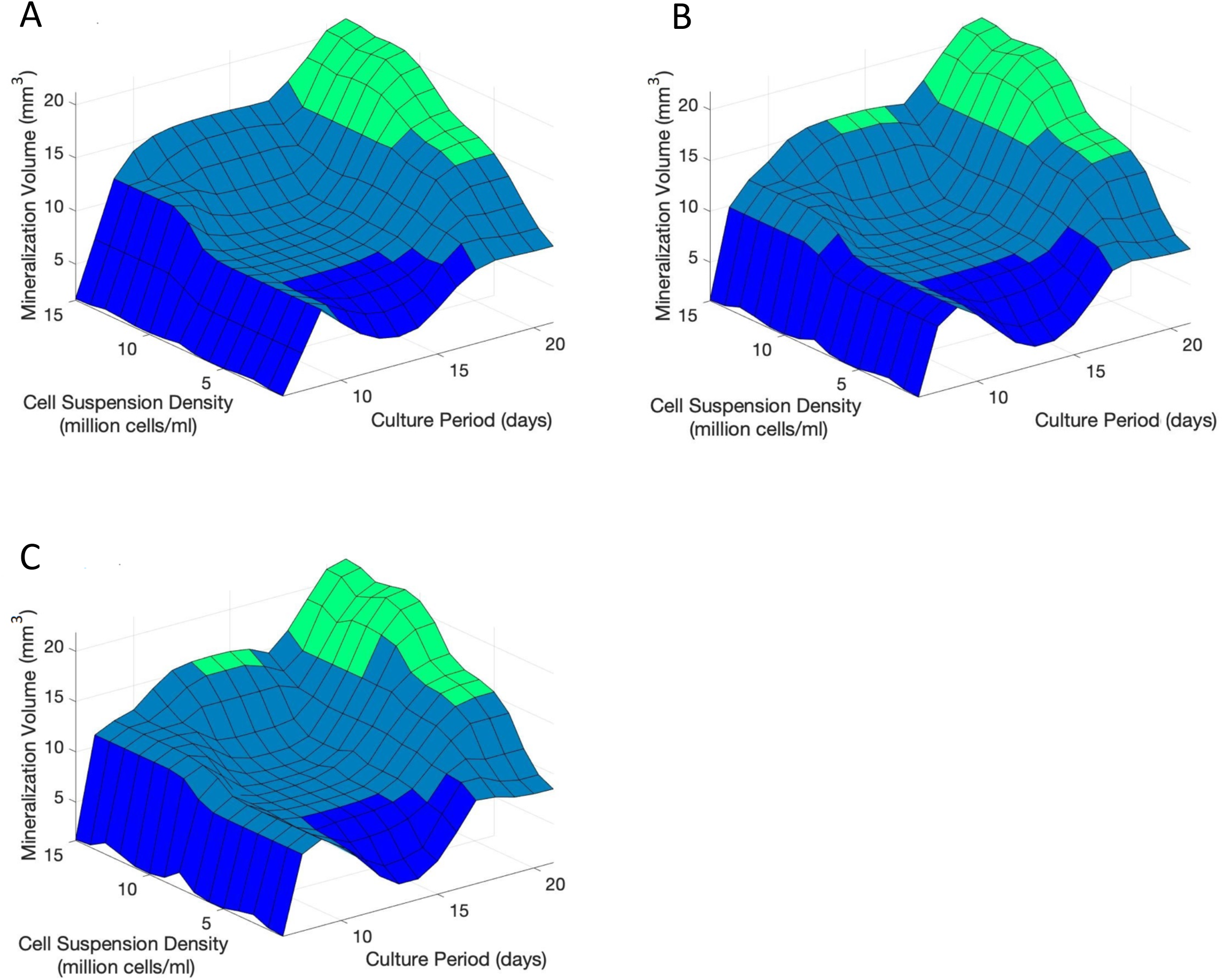
3D surfaces generated by implemented fuzzy rules. A) Type-1 fuzzy system. B) Type-2 fuzzy system with 20% uncertainty. C) Type-2 fuzzy system with 30% uncertainty.

### 4. Results

Type-1 and type-2 fuzzy systems were implemented as described above using input data from a previous experimental study [29]. Table 4 and Table 5 shows the T1 and T2 FS approximated values. In these tables, in addition to the experimental input data, the type-2 fuzzy system has an additional input, which is the uncertainty boundary (either 20% or 30%).This uncertainty boundary is used as an input to compare the mineralization volume output with different noise levels in the input data.

**Table 4.** Type-1 and Type-2 Fuzzy system approximation comparison

| Culture<br>Period<br>(days) | Cell<br>Suspension<br>Density<br>(million cells<br>/milliliter) | Mineralization Volume (mm<sup>3</sup>) |  |  |
| --- | --- | --- | --- | --- |
|  |  | Zero<br>Uncertainty<br>(T1 FS) | %20<br>Uncertainty<br>(T2 FS) | %30<br>Uncertainty<br>(T2 FS) |
| 7 | 3 | 1.84 | 1.7 | 1.92 |
| 7 | 8 | 2.04 | 2.1 | 2.73 |
| 7 | 11 | 1.75 | 1.32 | 1.31 |
| 7 | 14 | 1.78 | 1.38 | 1.40 |
| 14 | 3 | 7.22 | 6.72 | 6.75 |
| 14 | 8 | 8.41 | 8.37 | 8.46 |
| 14 | 11 | 9.43 | 8.89 | 8.91 |
| 14 | 14 | 14.5 | 15.17 | 15.13 |
| 21 | 3 | 11.53 | 11.14 | 11.15 |
| 21 | 8 | 17.07 | 17.28 | 17.22 |
| 21 | 11 | 20.76 | 21.51 | 21.56 |
| 21 | 14 | 21.13 | 21.6 | 21.7 |

**Table 5.** Accuracy of Type-1 and Type-2 (20% and 30% uncertainty) Fuzzy Systems

| Culture<br>Period<br>(days) | Cell<br>Suspension<br>Density<br>(million cells<br>/milliliter) | Mineralization Volume (mm<sup>3</sup>) |  |  |  |
| --- | --- | --- | --- | --- | --- |
|  |  | Experimental<br>Values<br>(Actual) | Zero<br>Uncertainty<br>(T1 FS) | 20%<br>Uncertainty<br>(T2 FS) | 30%<br>Uncertainty<br>(T2 FS) |
| 7 | 1.67 | 0.1 | 1.84 | 1.32 | 1.2 |
| 7 | 5 | 0.1 | 2.04 | 2.1 | 2.73 |
| 7 | 15 | 0.2 | 1.75 | 1.32 | 1.31 |
| 14 | 1.67 | 1 | 7.6 | 5.8 | 4.0 |
| 14 | 5 | 3 | 8.28 | 8.21 | 8.21 |
| 14 | 15 | 12 | 15.57 | 15.58 | 15.87 |
| 21 | 1.67 | 14 | 9.43 | 8.89 | 8.9 |
| 21 | 5 | 21 | 14.9 | 15.4 | 15.45 |
| 21 | 15 | 24 | 21.2 | 21.81 | 21.89 |
| **RMSE** |  | N/A | **3.6** | **3.79** | **3.5** |

First, we generated approximated values at each of the input value combinations available (Table 4). A graphical visualization of these values has been generated as a 3D surface, for the T1 FS (Fig. 3A), 20% T2 FS (Fig. 3B), and 30% T2 FS (Fig. 3C). Next, we compared the experimental output value (mineralization volume) to the approximated values in each of the three fuzzy systems (Table 5). Here, we calculated the root mean square error (RMSE) between the experimental and approximated values with same input values. The equation of the measurement is given as follow:

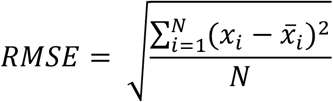

As compared to the T1 FS, we observed that the 20% T2 FS increased the overall error of the approximation (+5.3%). In contrast, we observed that the 30% T2 FS decreased the overall error of the approximation (−2.8%), relative to the T1 FS.

## 5. Discussion

Bioprinting research is a time-consuming and expensive process requiring the use of cells that may be difficult to source followed by a lengthy culture period. Finally, quantification of the suitability of the bioprint itself can be challenging. As a result, it is difficult to exhaustively optimize a bioprinted construct experimentally. Here, we have demonstrated that approximating bioprinting output parameters using fuzzy systems based on input variables is a viable approach to accelerate research, reduce experimental costs, and improve outcomes. Furthermore, our fuzzy system approach can be redesigned with additional input and output variables, qualitative results, or expert knowledge using linguistic rules.

In this study, we implement a fuzzy logic-based model using both type-1 and type-2 fuzzy systems to compare the results in handling the uncertainty associated with the bioprinting process. This uncertainty can arise from noisy input data or imperfect expert knowledge. Using experimental data, we have demonstrated that the implemented fuzzy logic can convert the discrete crisp input data to fuzzy sets to achieve a continuum data surface with high accuracy. Furthermore, we found that the 30% type-2 fuzzy system can accommodate more imprecision with higher accuracy, which may prove valuable for bioprinting optimization. In contrast, the 20% type-2 fuzzy system was unable to provide higher accuracy relative to the type-1 fuzzy system, so additional work may be necessary to determine the level of uncertainty specific to the biological process being optimized.

We have provided 3D surfaces generated by the fuzzy rules (e.g. Figure 3) as an intuitive tool to help researchers design new studies for experimental optimization of bioprinted constructs. For example, researchers should avoid performing new experiments in “flat” areas of the 3D model. In our study, the 3D surface illustrates a relatively flat area for moderate cell suspension densities and culture periods of around 14 days. Therefore, if the researcher wishes to maximize mineralization volume with a low number of cells (e.g. 5 x10^6^ cells/mL), they may prefer to increase their culture time to 21 days rather than attempt to triple their cell number in order to traverse the flat area of the surface. In total, implementation of this system may help researchers optimize their study design to eliminate unnecessary experimentation.

We also note that the 3D surfaces generated for the type-2 fuzzy systems (Figures 3B,C) are both qualitatively smoother than surface generated using the type-1 fuzzy system (Figure 3A). In particular, we observed a sharp edge around a culture period of 8 days. This smoothness is likely to result in higher accuracy in approximated values, as biological mathematical models ought to have a smooth transmission when increasing the input values.

Fuzzy systems are designed for decision-making, approximation, and optimization. A limitation of our method is the lack of a feedback system for incorporating new experimental results. Methods such as the Adaptive Neuro-Fuzzy Inference System (ANFIS) with Sugeno model give the fuzzy system a feedback feature to train and adjust the fuzzy membership functions based on new trained and tested data. Future studies should focus on implementing such a feature to further improve this approximation system for bioprinting.

## References

[1] A. Sedigh, A. R. Kachooei, P. K. Beredjiklian, A. R. Vaccaro, and M. Rivlin, “Safety and efficacy of casting during COVID-19 pandemic: A comparison of the mechanical properties of polymers used for 3D printing to conventional materials used for the generation of orthopaedic orthoses,” Arch. Bone Jt. Surg., vol. 8, no. SpecialIssue, 2020, doi: 10.22038/abjs.2020.44038.2204.

[2] A. Sedigh, M. H. Ebrahimzadeh, M. Zohoori, and A. Kachooei, “Cubitus Varus Corrective Osteotomy and Graft Fashioning Using Computer Simulated Bone Reconstruction and Custom-Made Cutting Guides,” Arch. Bone Jt. Surg., vol. 0, Nov. 2020, doi: 10.22038/abjs.2020.52457.2592.

[3] S. Derakhshanfar, R. Mbeleck, K. Xu, X. Zhang, W. Zhong, and M. Xing, “3D bioprinting for biomedical devices and tissue engineering: A review of recent trends and advances,” Bioactive Materials, vol. 3, no. 2. KeAi Communications Co., pp. 144–156, Jun. 01, 2018, doi: 10.1016/j.bioactmat.2017.11.008.

[4] S. V. Murphy and A. Atala, “3D bioprinting of tissues and organs,” Nature Biotechnology, vol. 32, no. 8. Nature Publishing Group, pp. 773–785, 2014, doi: 10.1038/nbt.2958.

[5] T. K. Koo and M. Y. Li, “A Guideline of Selecting and Reporting Intraclass Correlation Coefficients for Reliability Research.,” J. Chiropr. Med., vol. 15, no. 2, pp. 155–63, Jun. 2016, doi: 10.1016/j.jcm.2016.02.012.

[6] B. Webb and B. J. Doyle, “Parameter optimization for 3D bioprinting of hydrogels,” Bioprinting, vol. 8, pp. 8–12, Dec. 2017, doi: 10.1016/j.bprint.2017.09.001.

[7] A. Sedigh, J. E. Tulipan, M. R. Rivlin, and R. E. Tomlinson, “Utilizing Q-Learning to Generate 3D Vascular Networks for Bioprinting Bone,” bioRxiv, p. 2020.10.08.331611, Oct. 2020, doi: 10.1101/2020.10.08.331611.

[8] R. Suntornnond, E. Tan, J. An, and C. Chua, “A Mathematical Model on the Resolution of Extrusion Bioprinting for the Development of New Bioinks,” Materials (Basel)., vol. 9, no. 9, p. 756, Sep. 2016, doi: 10.3390/ma9090756.

[9] A. Torres and J. J. Nieto, “Fuzzy logic in medicine and bioinformatics,” Journal of Biomedicine and Biotechnology, vol. 2006. 2006, doi: 10.1155/JBB/2006/91908.

[10] J. Sun, J. M. Garibaldi, and C. Hodgman, “Parameter estimation using metaheuristics in systems biology: A comprehensive review,” IEEE/ACM Trans. Comput. Biol. Bioinforma., vol. 9, no. 1, pp. 185–202, 2012, doi: 10.1109/TCBB.2011.63.

[11] L. You, “Toward computational systems biology,” Cell Biochemistry and Biophysics, vol. 40, no. 2. Springer, pp. 167–184, Apr. 2004, doi: 10.1385/CBB:40:2:167.

[12] J. R. Banga, “Optimization in computational systems biology,” BMC Systems Biology, vol. 2, no. 1. BioMed Central, p. 47, May 28, 2008, doi: 10.1186/1752-0509-2-47.

[13] R. Küffner, T. Petri, L. Windhager, and R. Zimmer, “Petri Nets with Fuzzy Logic (PNFL): Reverse Engineering and Parametrization,” PLoS One, vol. 5, no. 9, p. e12807, Sep. 2010, doi: 10.1371/journal.pone.0012807.

[14] T. Z. Tan, G. S. Ng, and C. Quek, “A novel biologically and psychologically inspired fuzzy decision support system: Hierarchical complementary learning,” IEEE/ACM Trans. Comput. Biol. Bioinforma., vol. 5, no. 1, pp. 67–79, Jan. 2008, doi: 10.1109/TCBB.2007.1064.

[15] J. Bordon, M. Moskon, N. Zimic, and M. Mraz, “Fuzzy logic as a computational tool for quantitative modelling of biological systems with uncertain kinetic data,” IEEE/ACM Trans. Comput. Biol. Bioinforma., vol. 12, no. 5, pp. 1199–1205, Sep. 2015, doi: 10.1109/TCBB.2015.2424424.

[16] L. Z.-I. and control and undefined 1965, “Fuzzy sets,” Elsevier, Accessed: Nov. 05, 2020. [Online]. Available: https://www.sciencedirect.com/science/article/pii/S001999586590241X.

[17] S. Alayón, R. Robertson, S. K. Warfield, and J. Ruiz-Alzola, “A fuzzy system for helping medical diagnosis of malformations of cortical development,” J. Biomed. Inform., vol. 40, no. 3, pp. 221–235, Jun. 2007, doi: 10.1016/j.jbi.2006.11.002.

[18] E. M.-P. of the institution of electrical and undefined 1974, “Application of fuzzy algorithms for control of simple dynamic plant,” ieeexplore.ieee.org, Accessed: Nov. 05, 2020. [Online]. Available: https://ieeexplore.ieee.org/abstract/document/5250910/.

[19] N. N. Karnik, J. M. Mendel, and Q. Liang, “Type-2 fuzzy logic systems,” IEEE Trans. Fuzzy Syst., vol. 7, no. 6, pp. 643–658, Dec. 1999, doi: 10.1109/91.811231.

[20] Q. Liang and J. M. Mendel, “Interval type-2 fuzzy logic systems: Theory and design,” IEEE Trans. Fuzzy Syst., vol. 8, no. 5, pp. 535–550, Oct. 2000, doi: 10.1109/91.873577.

[21] N. N. Karnik and J. M. Mendel, “Centroid of a type-2 fuzzy set,” Inf. Sci. (Ny)., vol. 132, no. 1–4, pp. 195–220, Feb. 2001, doi: 10.1016/S0020-0255(01)00069-X.

[22] N. N. Karnik and J. M. Mendel, “Applications of type-2 fuzzy logic systems: handling the uncertainty associated with surveys,” in IEEE International Conference on Fuzzy Systems, 1999, vol. 3, doi: 10.1109/fuzzy.1999.790134.

[23] F. Baghbani, M. R. Akbarzadeh-T., and A. Akbarzadeh, “Indirect adaptive robust mixed H2/H∞ general type-2 fuzzy control of uncertain nonlinear systems,” Appl. Soft Comput. J., vol. 72, pp. 392–418, Nov. 2018, doi: 10.1016/j.asoc.2018.06.049.

[24] S. F. Toloue, M. R. Akbarzadeh, A. Akbarzadeh, and M. Jalaeian-F, “Position tracking of a 3-PSP parallel robot using dynamic growing interval type-2 fuzzy neural control,” Appl. Soft Comput. J., vol. 37, pp. 1–14, Dec. 2015, doi: 10.1016/j.asoc.2015.07.015.

[25] H. R. Hassanzadeh, M. R. Akbarzadeh-T, A. Akbarzadeh, and A. Rezaei, “An interval-valued fuzzy controller for complex dynamical systems with application to a 3-PSP parallel robot,” Fuzzy Sets Syst., vol. 235, pp. 83–100, Jan. 2014, doi: 10.1016/j.fss.2013.02.009.

[26] J. M. Mendel, “A comparison of three approaches for estimating (synthesizing) an interval type-2 fuzzy set model of a linguistic term for computing with words,” Granul. Comput., vol. 1, no. 1, pp. 59–69, Mar. 2016, doi: 10.1007/s41066-015-0009-7.

[27] N. N. Karnik and J. M. Mendel, “Type-2 fuzzy logic systems: type-reduction,” in Proceedings of the IEEE International Conference on Systems, Man and Cybernetics, 1998, vol. 2, pp. 2046–2051, doi: 10.1109/icsmc.1998.728199.

[28] K. Guney and N. Sarikaya, “COMPARISON OF MAMDANI AND SUGENO FUZZY INFERENCE SYSTEM MODELS FOR RESONANT FREQUENCY CALCULATION OF RECTANGULAR MICROSTRIP ANTENNAS,” 2009.

[29] J. Zhang et al., “Optimization of mechanical stiffness and cell density of 3D bioprinted cell-laden scaffolds improves extracellular matrix mineralization and cellular organization for bone tissue engineering,” Acta Biomater., vol. 114, pp. 307–322, Sep. 2020, doi: 10.1016/j.actbio.2020.07.016.

[30] B. M. Moreno-Cabezali and J. M. Fernandez-Crehuet, “Application of a fuzzy-logic based model for risk assessment in additive manufacturing R&D projects,” Comput. Ind. Eng., vol. 145, p. 106529, Jul. 2020, doi: 10.1016/j.cie.2020.106529.

[31] A. Sadollah, “Introductory Chapter: Which Membership Function is Appropriate in Fuzzy System?,” in Fuzzy Logic Based in Optimization Methods and Control Systems and its Applications, InTech, 2018.

